# Dispersal rate and riverine network connectivity shape the genetic diversity of freshwater amphipod metapopulations

**DOI:** 10.1101/2020.12.21.423851

**Authors:** Roman Alther, Emanuel A. Fronhofer, Florian Altermatt

## Abstract

Theory predicts that the distribution of genetic diversity in a landscape is strongly dependent on the connectivity of the metapopulation and the dispersal of individuals between patches. However, the influence of explicit spatial configurations such as dendritic landscapes on the genetic diversity and structure of metapopulations is still understudied, and theoretical corroborations of empirical patterns are largely lacking. Here, we used real-world microsatellite data and stochastic simulations of two metapopulations of freshwater amphipods in a 28,000 km^2^ riverine network to study the influence of spatial connectivity and dispersal strategies on their spatial genetic diversity and structure. We found a significant imprint of the riverine network connectivity on the genetic diversity of both amphipod species. Data from 95 sites showed that allelic richness and observed heterozygosity significantly increased towards more central nodes of the network. In simulation models, dispersal rate was suggested to be the key factor explaining the empirically observed distribution of genetic diversity. Contrary to often-claimed expectations, however, the relevance of directionality of dispersal was only minor. Surprisingly, also the consideration of site-specific carrying capacities, for example by assuming a direct dependency of population size with local river size, substantially decreased the model fit to empirical data. This highlights that directional dispersal and the spatial arrangement of population sizes may have a smaller relevance in shaping population genetic diversity of riverine organisms than previously thought, and that dispersal along the river network is the single-most important determinant of population genetic diversity.

## Introduction

The genetic diversity of populations is shaped by gene flow, selection, and genetic drift (Hartl & Clark, 2006; Manel et al., 2003). These processes are interacting with ecological processes, determining the organisms’ demography, population size and population dynamics. Understanding both ecological and genetic processes affecting natural populations is thus central to the understanding of patterns and dynamics of biological diversity and for implementing appropriate conservation strategies (Balkenhol et al., 2016; Lande, 1988), especially in the context of habitat fragmentation.

Extensive theoretical and empirical work highlights that dispersal has a pronounced effect on the genetic diversity and effective size of the population (Bowler & Benton, 2005; Clobert et al., 2012) and the community composition of populations (Vellend, 2016). Dispersal is defined by the movement of organisms with potential consequences for gene flow (Ronce, 2007), and is especially relevant in spatially structured landscapes (Gilpin & Hanski, 1991; Hanski & Simberloff, 1997). Thereby, multiple aspects of dispersal have been proposed to explicitly modulate dispersal, such as the rate of dispersal, the specific spatial dispersal trajectories (i.e., landscape connectivity), or local population sizes (Hanski & Gaggiotti, 2004). These need to be considered to understand the overall effects of dispersal on the genetic diversity of natural populations. Understanding the importance of different aspects of dispersal for natural populations is empirically challenging because both the processes influencing individual dispersal and its population genetic consequences need to be explored simultaneously.

The study of dispersal has a long tradition in both landscape ecology and metapopulation ecology, respectively, using slightly different tools and perspectives (Clobert et al., 2012; Hanski & Gaggiotti, 2004; Leibold et al., 2004; Vellend, 2016). Ideally, approaches combine measures of genetic and landscape connectivity, linking physical connectivity to genetics (Manel & Holderegger, 2013). The metapopulation approach provides the means of including connectivity more explicitly (Hanski & Gaggiotti, 2004). Studies about the influence of landscape connectivity on the genetic structure of populations mostly study lattice-like landscapes (2D), such as grasslands or forests (Dyer et al., 2012; Fortuna et al., 2009; Rozenfeld et al., 2008), using least-cost path methods with landscape resistance to integrate their spatial complexity (Adriaensen et al., 2003; Pinto & Keitt, 2009; Wang et al., 2009). However, this may not be generalized to the spatial structure of all ecosystems, and dispersal of organisms may be differently and stronger confined in spatially more complexly structured ecosystems.

Riverine systems are a prominent example thereof. Their connectivity is highly characteristic, can be explicitly quantified, and follows a universal dendritic network structure. These systems are formed by geological processes leading to general topological patterns (Altermatt, 2013; Carraro et al., 2020; Rodríguez-Iturbe & Rinaldo, 1997). Ecological consequences of the spatial configuration in such networks are well-studied, and encompass effects on species richness, on beta-diversity, as well as on population sizes (Altermatt et al., 2013; Carrara et al., 2012; Henriques-Silva et al., 2019; Muneepeerakul et al., 2008; Tonkin et al., 2018). In contrast, the evolutionary consequences of the network structure on the intraspecific genetic diversity are less well understood, even though one could apply the same approaches to study them. Comparative studies focusing on the effect of riverine network structures on intraspecific genetic diversity are still rare (Blanchet et al., 2020; Brauer et al., 2018; Fourtune et al., 2016), but generally show an increase in diversity in more downstream parts of the network (“downstream increase in intraspecific genetic diversity” (DIGD); Paz-Vinas et al., 2015). Importantly, these studies highlight that various processes can lead to such empirically observed patterns (Blanchet et al., 2020). In parallel, theoretical models that address the effect of spatial connectivity of riverine networks on genetic variation (Morrissey & de Kerckhove, 2009; Paz-Vinas et al., 2015; Paz-Vinas & Blanchet, 2015), on evolution of dispersal (Henriques-Silva et al., 2015), and emergence of neutral genetic structure (Fronhofer & Altermatt, 2017; Stokes & Perron, 2020; Thomaz et al., 2016) have demonstrated that dispersal along riverine networks has a direct imprint on the genetic structure and diversity of the inhabiting organisms. While these theoretical models provide direct testable predictions, a direct comparison of empirical data and predictions of theoretical models has been largely lacking.

Here, we studied the influence of connectivity in a real-world riverine network on the genetic diversity of freshwater amphipods (crustaceans), by combining large-scale empirical data on their population genetic diversity and structure with a simulation model making analogue predictions of their population genetic diversity using a graph theoretic approach. Graph theory has not yet been widely used in landscape genetics although it allows concise presentation of spatial configuration of natural populations (Dyer & Nason, 2004; Fortuna et al., 2009; Garroway et al., 2008; Manel & Holderegger, 2013). Based on previous work (Altermatt & Fronhofer, 2018; Blanchet et al., 2020; Fronhofer & Altermatt, 2017; Muneepeerakul et al., 2007), we wanted to gain a better understanding of the relative importance of different ecologically relevant aspects of dispersal on shaping the genetic structure of populations. In particular, we expected allelic richness and observed heterozygosity to be higher in more central nodes of the riverine network, (Paz-Vinas et al., 2015; Ritland, 1989). This is caused by a strong signal of dispersal rate, upstream movement probability, and habitat carrying capacity leading to central nodes receiving more migrants and sustain larger populations (Blanchet et al., 2020; compare Paz-Vinas et al., 2015). We addressed this with microsatellite data from 3,319 amphipod individuals collected from 95 sites across a riverine network covering more than 28,000 km^2^ and compared it to the output of stochastic simulation models examining alternative parameter combinations influencing dispersal, but conducted on the identical riverine network structure.

## Methods

### Study system

*Gammarus fossarum* (Koch) is a common and wide-spread freshwater amphipod species complex (Crustacea, Amphipoda), predominantly found in smaller streams and distributed throughout Central Europe (Karaman & Pinkster, 1977; Wattier et al., 2020; Weiss et al., 2014) and adjacent biogeographic regions. As a major decomposer, it has an important role in aquatic food webs (Hieber & Gessner, 2002; Little & Altermatt, 2018). The species complex contains a high cryptic diversity, with several to dozens of species being reported, but not yet formally described (Müller, 2000; Wattier et al., 2020; Weiss et al., 2014). In Switzerland, two of those cryptic lineages are widely distributed (Altermatt et al., 2014, 2019; Westram et al., 2011, 2013). These lineages are reproductively isolated, and are “good” species that diverged ∼15 Mio years ago (Wattier et al., 2020), herein referred to as *Gammarus fossarum* type A (*G. fossarum* A) and *Gammarus fossarum* type B (*G. fossarum* B, both sensu Müller (2000)). While reproductively isolated, their distributional range and their ecological functions have a substantial overlap (Eisenring et al., 2016; Müller et al., 2000; Wattier et al., 2020). This allows treating them as two biological replicates of species to study effects of spatial network structure on the genetic structure of (meta)populations (Altermatt et al., 2019; Eisenring et al., 2016).

### Genetic data collection

We conducted the study in the river Rhine drainage within Switzerland, which encompasses about 28,000 km^2^ of its headwater area. We sampled *Gammarus fossarum* amphipods from 281 sites evenly and representatively spaced across the river Rhine headwaters between 2007 and 2015 by a kicknet approach. We morphologically identified all to the species-complex level (Altermatt et al., 2019). We further genotyped a subset of individuals of the *G. fossarum* complex using microsatellites (Westram et al., 2013), conventional 16S sequencing, or SNP pyrosequencing (Westram et al., 2011). In this study, we used only the microsatellite data for subsequent population genetic analyses, whereas we used 16S and SNP data to distinguish between *Gammarus fossarum* type A and type B.

We extracted DNA for microsatellite analyses from complete individuals or their heads using a HotSHOT approach (Montero-Pau et al., 2008). We amplified fragments using multiplex amplifications with the QIAGEN Multiplex PCR Kit chemicals. We used ten different microsatellite markers (gf08, gf10, gf13, gf18, gf, 19, gf21, gf22, gf24, gf27, gf28 sensu Westram et al. (2010)), specifically designed for *G. fossarum* and previously established in several studies (Altermatt et al., 2014; Westram et al., 2013). We used primers in different concentrations (see table 1 in Westram et al., 2010) in reaction volumes of 12.5 μl, with 6.25 μl of PCR Master Mix, 1.25 μl Q solution and 1 μl DNA template. The PCR consisted of an initial denaturation step at 95 °C (15 min), 35 cycles at 94 °C (30 s), 60 °C (90 s), 72 °C (60 s), and a final elongation step at 60 °C (30 min). We diluted the resulting amplicons (1:20) and combined them with size standard (GeneScan(tm) 500 LIZ(tm), Applied Biosystems). We sequenced fragments on an Applied Biosystems 3730xl DNA Analyzer at the Genomic Diversity Center of ETH Zurich, Switzerland. We analyzed and manually edited the electropherograms using SoftGenetics GeneMarker software (v. 1.80). In total, we genotyped 3,577 individuals. We used the microsatellite data to quantify genetic diversity and differentiation within these two species. For the detailed molecular procedure on DNA extraction, microsatellite sequencing and microsatellite interpretation, see Westram et al. (2010, 2013), in which some of the individuals used here have already been analyzed for different purposes.

**Table 1.** Chosen simulation parameters, explored values and their biological meaning.

| Parameter | Values | Meaning |
| --- | --- | --- |
| *Varying parameters* |  |  |
| $d$ | 0.001; 0.01; 0.1 | Dispersal rate |
| $W$ | 0; 0.5; 1 | Upstream movement probability |
| $K$ | fixed=1000; scaled=[2; 8,992] | Carrying capacity |
| *Fixed parameters* |  |  |
| $\mu$ | 0.0001 | Mutation rate neutral alleles |
| $\lambda_0$ | 2 | Fecundity |
| $m$ | 0 | Dispersal mortality |

### Spatial data preparation

The spatial riverine network used for the subsequent analysis represents a restricted version of the full Rhine network within Switzerland. We constructed a digital representation of the riverine network (Figure S1), based on a graph theory approach and following topological connectivity along the river lines. The riverine network is based on a 2 km^2^ subcatchment representation of streams and rivers of Switzerland (BAFU, 2012). Specifically, we interpreted these subcatchments as being nodes within the network and stream flow direction being directed vertices between these nodes. Based on the coordinates of the outlet site of each subcatchment (or the centroid coordinates if headwater subcatchment), we constructed the riverine network connecting the outlet coordinates to each other. Distances from one subcatchment outlet to the adjacent downstream subcatchment outlet can be approximated as Euclidean distances on a small-scale basis, and were included as vertex weights. Additionally, the graph object also contained information on the summed upstream catchment area for each subcatchment. The detailed methods of how we prepared the extensive graph object are described in Alther & Altermatt (2018).

We then restricted the analysis to the part of the riverine network that is actually inhabitable by either one or both of the studied amphipod species, based on an empirically validated cropping of the network. We used a database on amphipod occurrences in Switzerland with >2,000 sites covered (Altermatt et al., 2019) to distinguish between nodes containing *G. fossarum* from unoccupied nodes. After preparation of the initial complete Rhine riverine network, we selected nodes (subcatchments) that contain one or both species of the *G. fossarum* complex and all their spatially interconnecting nodes (Figure S1). This resulted in the truncated riverine network that is empirically validated to be accessible and inhabitable to *G. fossarum*, containing a total of 2,401 nodes (referred to as *G. fossarum* network). The corresponding graph object is available on GitHub (Gfos_Network.RData: DOI 10.5281/zenodo.4321240).

We subsequently mapped our microsatellite data from *G. fossarum* complex sites of Switzerland to the nodes of the prepared graph using ArcGIS 10.5.1 (ESRI Inc., Redlands, California, USA). We refer to sites as nodes hereafter. Removing nodes that had less than 15 individuals successfully genotyped resulted in 95 nodes for subsequent analysis, harboring 3,319 individuals. The corresponding microsatellite data are available on GitHub (Gammarus_all_microsat_data_Rhein_2018.txt; DOI 10.5281/zenodo.4321240). Data were available for 67 nodes for *G. fossarum* type A (2,257 individuals) and 33 nodes for *G. fossarum* type B (1,062 individuals), with five nodes having data on both (Figure S1). This preparation step resulted in a vector containing the corresponding node IDs where microsatellite data of either one or both species of the *G. fossarum* complex were available.

### Stochastic simulation

To compare the empirical data to simulated data, we used a discrete-time and stochastic individual-based simulation (adapted from Fronhofer et al., 2013, 2014; Fronhofer & Altermatt, 2017). The model is analogous to the one used by Fronhofer & Altermatt (2017) and a detailed model description can be found there. In brief, we model a metapopulation of amphipods where we assume that local populations of amphipods compete for local and limited resources, which is captured by a Beverton-Holt density-regulation function (Beverton & Holt, 1957). Individuals are diploid and reproduce sexually (sex ratio: 0.5). Individuals perform natal, nearest-neighbor dispersal, which is governed by a dispersal rate (*d*), and by the connectivity matrix of the metapopulation that is identical to the one derived for the empirical data (*G. fossarum* network). Most of the parameters of the simulation were fixed (see Table 1) but informed by the study system or the empirical methods used. We assume ten neutral, diploid loci that can take any of 100 different values as alleles to explore genetic diversity. The mutation rate of those alleles is set to 0.0001 and the mean carrying capacity equals 1,000 individuals.

We subsequently explored the full-orthogonal parameter space along three free parameters. This included 1) three different dispersal rates (*d* = 0.001, 0.01, or 0.1); 2) three different upstream movement probabilities (*W*); and 3) two scenarios for the distribution of carrying capacities. Upstream movement probability described the effects of downstream water flow, where there was either no upstream movement (*W* = 0), where upstream and downstream movements were equally likely (*W* = 1), or where downstream movements was twice as likely as upstream movement (*W* = 0.5). Carrying capacity (*K*) was either identical for all nodes (*K* = 1000 per node), or it scaled with the square-root of the total catchment area as described by Rodriguez-Iturbe & Rinaldo (1997), such that the highest carrying capacity corresponded to the most downstream node while keeping the total metapopulation size constant (2,401 nodes × 1,000 individuals). All simulations were run with ten replicates, and for 10,000 generations each, which is sufficient to reach (quasi-)equilibrium. All population genetic analyses where performed on the individuals of the last generation (*t* = 10,000) in analogy to the empirical data analysis. The explored parameter space is detailed in Table 1. The simulation code is available on GitHub (pop_gen_gammaridae_v0.cpp: DOI 10.5281/zenodo.4321240).

### Statistical analyses

For the spatial data (explanatory variables), we calculated a series of network metrics in order to identify the influence of network topology based on the *G. fossarum* network containing 2,401 nodes and the subset of 95 nodes with microsatellite data available. The calculations were done in R 3.6.1 (R Core Team, 2019) with the package igraph 1.2.4.2 (Csárdi & Nepusz, 2006). The network metrics for single nodes were upstream distance from the outlet node, total upstream catchment area, directed and undirected betweenness centrality, directed and undirected closeness centrality, and degree centrality. Directed and undirected measures correspond to either considering flow direction, or ignoring it. The upstream distance corresponds to the instream distance from the outlet node of the riverine network near Basel, where the river Rhine continues to France and Germany and gets into a different biogeographic zone, thereby naturally separating the catchments considered here. The closeness centrality corresponds to the reciprocal of the sum of the distances between a node and all other nodes in the riverine network. We standardized the closeness centrality (*c)* for analysis using the following approach: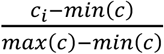. In biological terms, all these network metrics capture the connectivity of single populations to the other populations, with higher values translating to reduced connectivity.

For the genetic data (response variables), we calculated mean allelic richness and observed heterozygosity for *G. fossarum* type A and *G. fossarum* type B using the package hierfstat 0.04-22 (Goudet, 2005) within R 3.6.1 (R Core Team, 2019). We calculated these measures for both the empirical data and the simulated data.

Prior to modelling the empirical genetic response variables, we excluded highly correlated response variables (Kendall’s Tau > 0.8), specifically undirected betweenness centrality, degree centrality, and directed closeness centrality. We first modeled the genetic response variables separately using Generalized Linear Models (GLM) including all network metrics (upstream distance, total catchment area, directed betweenness centrality, undirected closeness centrality) and species as factor with all higher-level interaction terms. For mean allelic richness of populations, we applied Gamma GLMs, because all values are continuous but have a lower bound at zero. For mean observed heterozygosity, we applied quasibinomial GLMs, because the response values theoretically range from zero to one. For the Gamma GLMs we applied backward stepwise selection (function step()) using AIC scores in order to check if the interaction terms should remain in the model. Model selection for quasibinomial GLMs was done applying a backward model selection strategy by single term deletions of the explanatory variables and consecutively removing the variable with the highest P-value one at a time, updating the GLM and repeating the process. This initial exploration showed that models with higher-level interactions performed worse than models without interactions based on AIC (data not shown). Hence, we ran each a GLM for mean allelic richness and for mean observed heterozygosity including all explanatory variables without interaction terms. We additionally conducted separate GLMs for all explanatory variables individually for a qualitative comparison. GLMs were fitted using the glm() function. Figures were plotted using the fitted values retrieved from the separate GLMs with only one explanatory variable each, using the predict() function. F-test statistics for Gamma and quasibinomial GLMs were calculated using anova() with corresponding null models. R^2^_V_ as coefficient for determination (Zhang, 2017) was calculated using the rsq() function from the package rsq 2.0.

To assess which parameter combination for the simulations best fit the observed data, we compared simulated to empirical population genetic variables (mean allelic richness, mean observed heterozygosity). If model results and empirical data were identical, they would lie on the 1:1 diagonal line when plotting empirical vs. simulated data from identical nodes. Therefore, we use the perpendicular offset (distance) from the 1:1 diagonal to calculate goodness-of-fit measures. Unlike a conventional correlation, this also assesses the fits to both the range (intercept) and the explicit arrangement (slope) of response variables. We used the sum of perpendicular offsets (SPO) as well as the median of the perpendicular offsets (MPO) as goodness-of-fit measures. The SPO takes into account the overall spread of simulated values from their empirical counterpart, where a larger SPO indicates a poorer fit.

Considering the median using MPO partially takes into account outliers of individual nodes. An MPO closer to zero indicates that most of the simulated values fell close to the empirical counterpart. Since the perpendicular offset does not take into account if the offset is above or below the vertical (1:1) line, we additionally computed the directed median of the perpendicular offset (DMPO). To assess which specific parameter value for each of the varying parameters (dispersal rate, upstream movement probability, scaling of carrying capacity) generally best fitted to the observed data, we compared simulations with a specific parameter value to all corresponding simulations with the remaining parameter values of the same type. Specifically, we subtracted their goodness-of-fit measures (both, SPO and MPO) and assessed the sign (positive or negative). For example, the SPO for mean allelic richness with *d* = 0.001, *W* = 0, *K* = 0 was subtracted from the SPO for mean allelic richness with *d* = 0.01; *W* = 0; *K* = 0. If the simulation fit was higher for *d* = 0.001, this subtraction would result in a negative sign. Repeating this across all simulation combinations and both response variables (mean allelic richness, mean observed heterozygosity) resulted in a fraction of comparisons with a negative sign. If this fraction was higher than 0.5, the former parameter value was considered superior.

Finally, we used smoothed response variables to remove small-scale patterns driven by single sampling points and consequently focusing on a match with large-scale patterns. We compared simulated and empirical spatial patterns of population genetic diversity using local polynomial regressions fits for both types of data. We fitted local polynomial regressions using the loess() function with α = 0.5. We calculated the SPO, the MPO, and the DMPO using our own functions, included in the analysis script available on GitHub (PopGenNet20201210.R: DOI 10.5281/zenodo.4321240). All calculations were done in R ver. 3.6.1 (R Core Team, 2019).

## Results

### Population genetics of Gammarus fossarum *complex*

Empirically assessed mean allelic richness over all ten loci as a measure of local genetic diversity ranged from 1.6 to 5.3 (mean: 3.5; median: 3.5; SD: 0.8) for *G. fossarum* type A, and from 2.5 to 5.1 (mean: 3.4; median: 3.1; SD: 0.7) for *G. fossarum* type B. Both species showed clear geographic population genetic structuring in mean allelic richness (Figure 1a,b).

**Figure 1.**
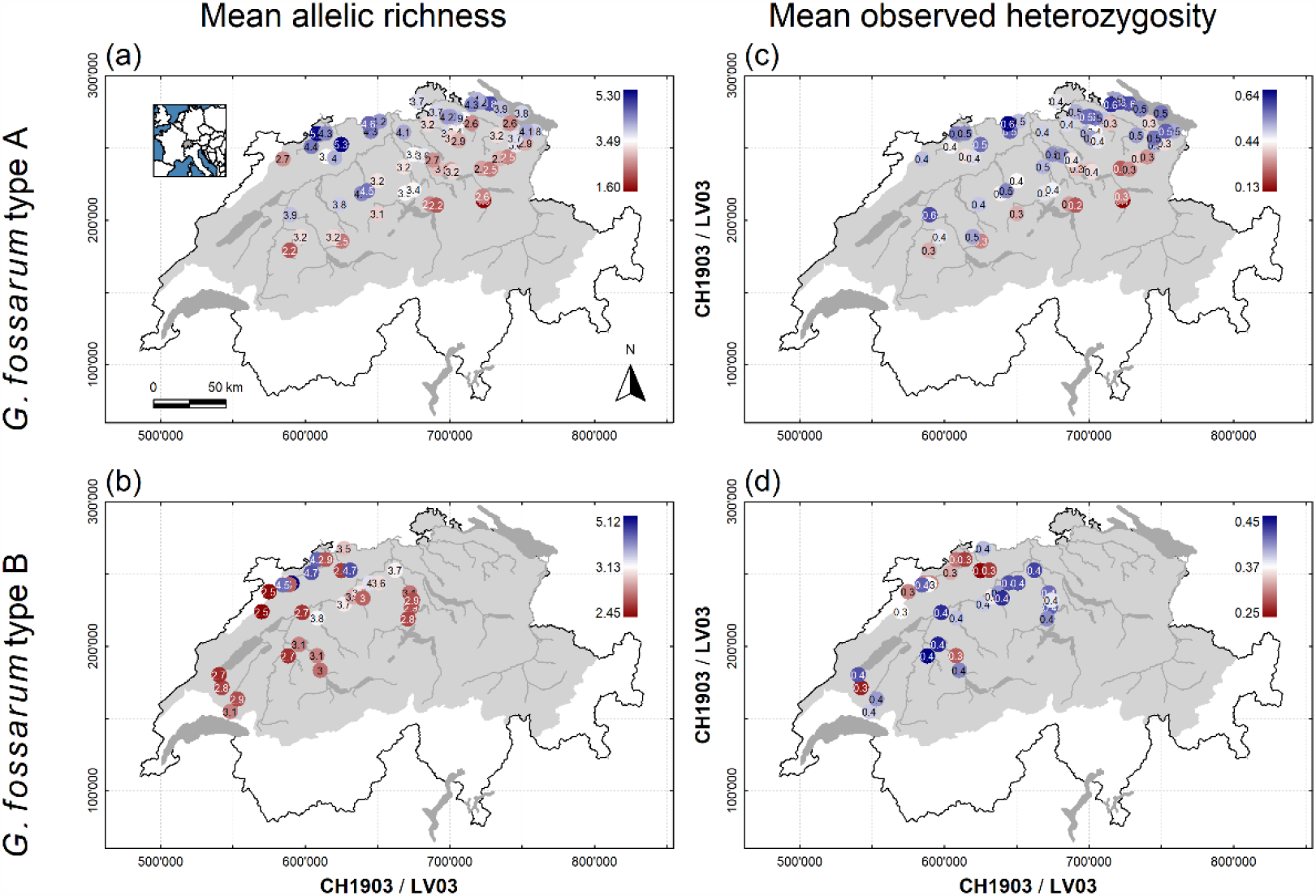
Empirically assessed mean allelic richness (a, b) and mean observed heterozygosity (c, d) of the two cryptic amphipod species *Gammarus fossarum* type A (a, c) and *G. fossarum* type B (b, d) in the river Rhine network in Switzerland. The river Rhine catchment is highlighted by the grey contour area, with the major river and lakes indicated. Both species of the *Gammarus fossarum* complex are widely distributed at elevations below 1000 m a.s.l., with type A being more common in the North Eastern part of the catchment, and type B more common in the Western part. Geodata source: Federal Office of Topography & Federal Office for the Environment.

Mean allelic richness was best explained by upstream distance in combination with total catchment area and species (Figure 2a; Gamma GLM; F_3,96_ = 13.3; *P* < 0.001; R^2^_V_ = 0.33). GLMs with a single explanatory network variable and species as factor showed that mean allelic richness significantly decreased with upstream distance from the outlet node within the riverine network in both species of the *Gammarus fossarum* complex; Gamma GLM; F_2,97_ = 16.2; *P* < 0.001; R^2^_V_ = 0.28). This translates to higher allelic richness in more central and better-connected nodes of the network. Allelic richness also significantly increased with standardized undirected closeness centrality when excluding other network metrics (Figure 2b; Gamma GLM; F_1,98_ = 4.5; *P* = 0.036; R^2^_V_ = 0.02), although the model fit was low. Mean allelic richness did not significantly increase with total catchment area (Gamma GLM; F_1,98_ = 3.5; *P* = 0.065; R^2^_V_ = 0.04). Directed betweenness centrality did not explain mean allelic richness (Gamma GLM; F_1,98_ = 3.4; *P* = 0.069; R^2^_V_ = 0.03).

**Figure 2.**
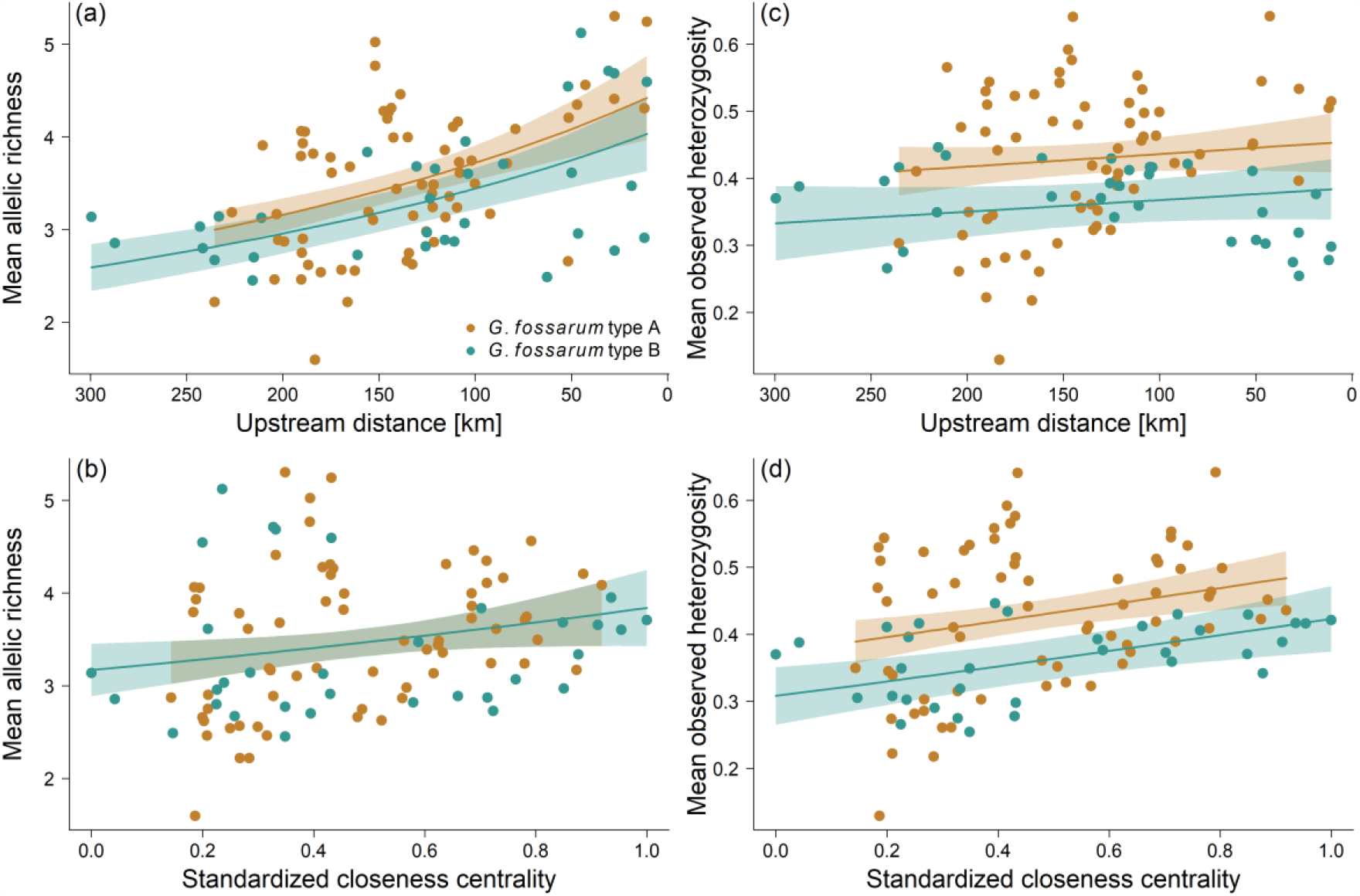
Empirically assessed mean allelic richness and mean observed heterozygosity of both species of the *Gammarus fossarum* complex (type A: orange points, type B: cyan points) with respect to different river network metrics. Raw data points as well as model fits of Generalized Linear Models (solid lines) and their 95% confidence intervals (shading) are given. Mean allelic richness as a function of (a) upstream distance from the outlet node within the riverine network and as a function of (b) standardized undirected closeness centrality. Mean observed heterozygosity as a function of (c) upstream distance from the outlet node within the riverine network and as a function of (d) standardized undirected closeness centrality.

Empirically assessed mean observed heterozygosity ranged from 0.13 to 0.64 (mean: 0.43; median: 0.44; SD: 0.11) for *G. fossarum* type A, and from 0.25 to 0.45 (mean: 0.36; median: 0.37; SD: 0.06) for *G. fossarum* type B. Geographic structuring of mean observed heterozygosity was only apparent in *G. fossarum* type A (Figure 1c) but not in *G. fossarum* type B (Figure 1d). For mean allelic richness the most parsimonious GLM relied on upstream distance and total catchment area as explanatory variables, while mean observed heterozygosity was best explained by undirected closeness centrality, directed betweenness centrality, and species (Figure 2d; Quasibinomial GLM; F_3,96_ = 9.4; *P* < 0.001; R^2^_V_ = 0.22). Observed heterozygosity was higher in more central and better-connected nodes of the network, hence it increased with increasing closeness centrality and higher betweenness centrality. The result from the GLM with only upstream distance as explanatory variable (the best fitting model for mean allelic richness) showed that heterozygosity increased closer towards the outlet node (decreasing upstream distance) for both species of the *G. fossarum* complex (Figure 2c; Quasibinomial GLM; F_2,97_ = 6.3; *P* = 0.003; pseudo-R^2^_V_ = 0.11). Mean observed heterozygosity also significantly increased with total catchment area in the GLM with only total catchment area and species included (Gamma GLM; F_2,97_ = 7.3; *P* = 0.001; R^2^_V_ = 0.12).

### Simulation – data comparison

The stochastic simulations resulted in highly differentiated spatial patterns of population genetic diversity depending on the set of parameter values used (Figure 3, Figure S2). Of the three different parameters considered (dispersal rate, upstream movement probability, scaling of carrying capacity), dispersal rate was generally showing the strongest effect on the response variable with respect to the parameter space covered (Figure S3). Using different upstream movement probabilities in the stochastic simulations resulted in smaller effects on the genetic diversity (shift of overall median of perpendicular offsets), but unidirectional movement (*W* = 0) generally resulted in better model fits (Figure S4). Scaling the carrying capacity (*K* = 1) consistently worsened the model fits and showed a smaller effect on the response variable compared to dispersal rate (Figure S5).

**Figure 3.**
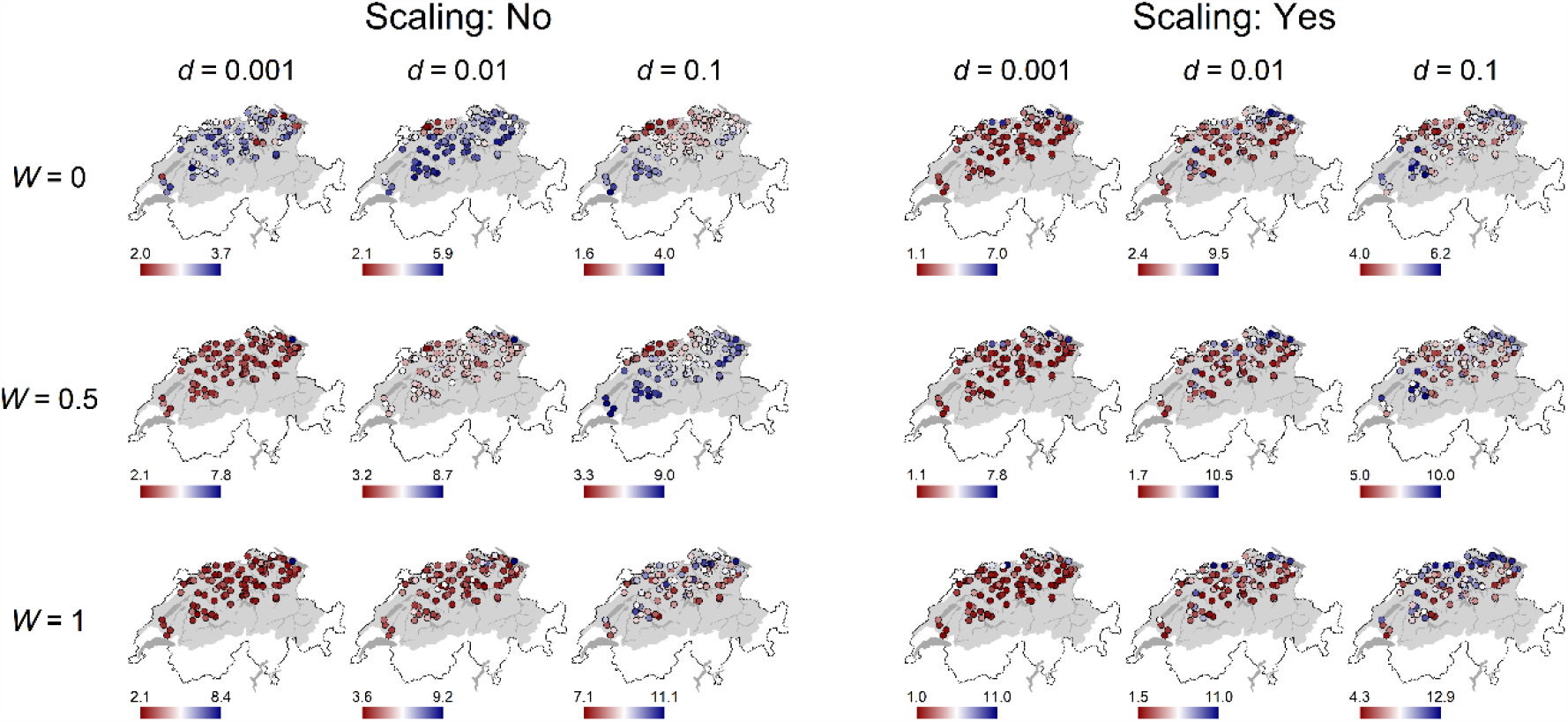
Maps depicting the predicted mean allelic richness for all 18 stochastic simulation scenarios show different spatial structuring along the Rhine riverine network of Switzerland. The gradient legends show mean allelic richness. Their scale is individually adjusted in each map for the best representation of spatial structuring. The corresponding figure for observed heterozygosity is given in Figure S2. Geodata source: Federal Office of Topography & Federal Office for the Environment.

Comparing the simulation outputs to empirical data showed that simulations based on low dispersal rates (*d* = 0.001) outperformed the corresponding ones with higher dispersal rates (*d* = 0.01 or *d* = 0.1) according to both goodness-of-fit measures (smaller SPO and smaller MPO) in 83% of the cases (80 comparisons out of 96). Simulations with no upstream dispersal (*W* = 0) outperformed simulations allowing some level of upstream dispersal (*W* = 0.5 and *W* = 1) in 77% of the cases (74 comparisons out of 96). Simulations with no scaling of carrying capacity (*K* = 0) outperformed their counterparts with scaling in 65% of the cases (47 comparisons out of 72). The best fitting simulations for allelic richness according to SPO for both species were based on low dispersal rates (*d* = 0.001), no upstream movement (*W* = 0), and no scaling of carrying capacity (Figure 4 and Figure S6, Table S8). For observed heterozygosity we also found that the best fitting simulation according to SPO was based on low dispersal rates (*d* = 0.001), no upstream movement (*W* = 0), and no scaling of carrying capacity in *G. fossarum* type A. In *G. fossarum* type B, the best simulation fit required high dispersal (*d* = 0.1), no upstream movement (*W* = 0), and no scaling of carrying capacity (Figure 5 and Figure S7, Table S9). When considering outliers by comparing MPO, the best fitting simulations for allelic richness and observed heterozygosity for *G. fossarum* type A were based on low dispersal rates (*d* = 0.001), no upstream movement (*W* = 0), and no scaling of carrying capacity (Figure 4 and Table S10, S11). With *G. fossarum* type B, the best fitting simulation for allelic richness required high dispersal (*d* = 0.1), no upstream movement (*W* = 0), and no scaling of carrying capacity (Figure 4 and Table S10). For observed heterozygosity, it required low dispersal (*d* = 0.001) and no scaling of carrying capacity as in *G. fossarum* type A, but coupled with upstream movement (*W* = 1) (Figure 5 and Table S11). The directed perpendicular offsets (DMPO) showed that simulations with low dispersal (*d* = 0.001) mostly fell within the range of observed empirical data. Simulations with higher dispersal rates (*d* = 0.01 or *d* = 0.1) consequently overestimated both measures of population genetic diversity (Figure S12 and S13), except when a high dispersal rate (*d* = 0.1) was coupled with no upstream dispersal (*W* = 0) and no scaling of carrying capacity.

**Figure 4.**
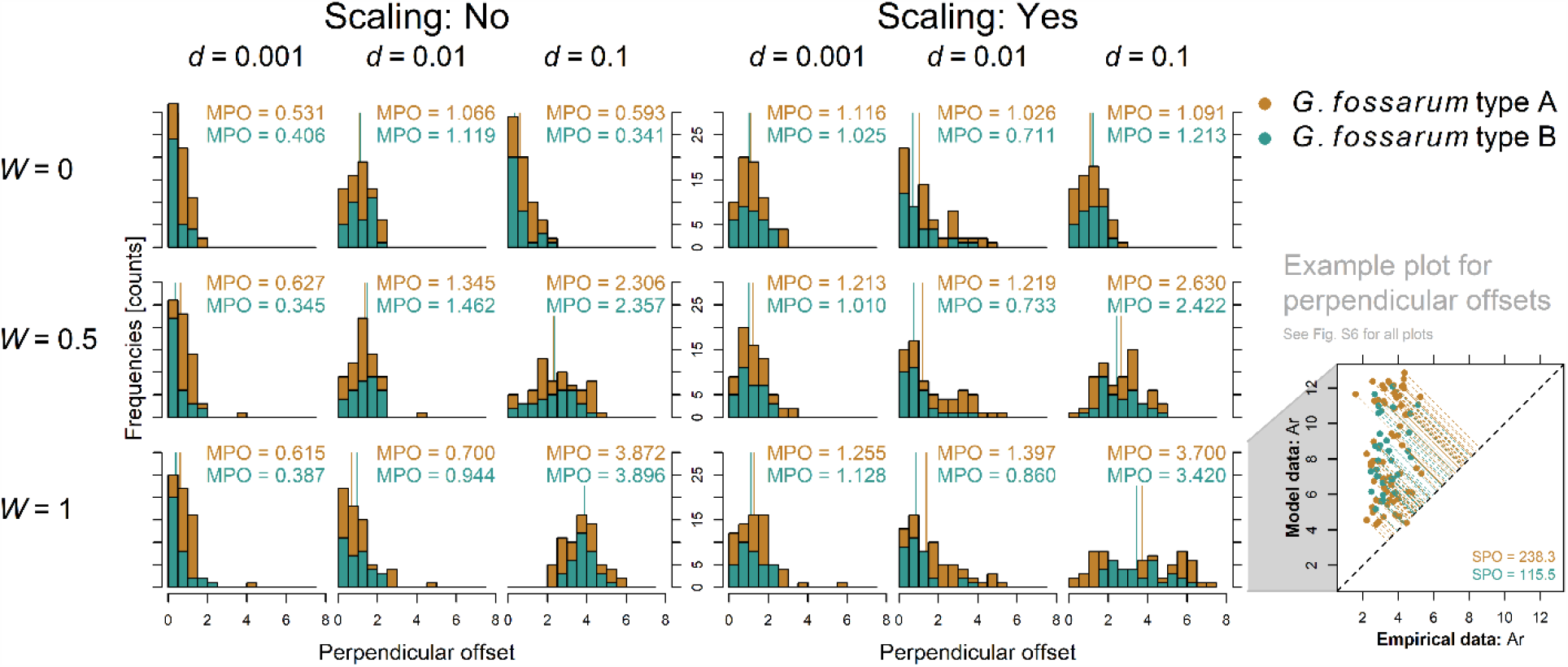
Histograms and medians of the perpendicular offsets (MPO) between all 18 stochastic simulation scenarios and the empirically assessed mean allelic richness values for both species of the *Gammarus fossarum* complex (type A: orange color, type B: cyan color). The more left-skewed a distribution, the better the fit of simulated values to empirical data. The example plot on the right hand side illustrates the concept of perpendicular offsets (compare to Figure S6 for all 18 scenarios).

**Figure 5.**
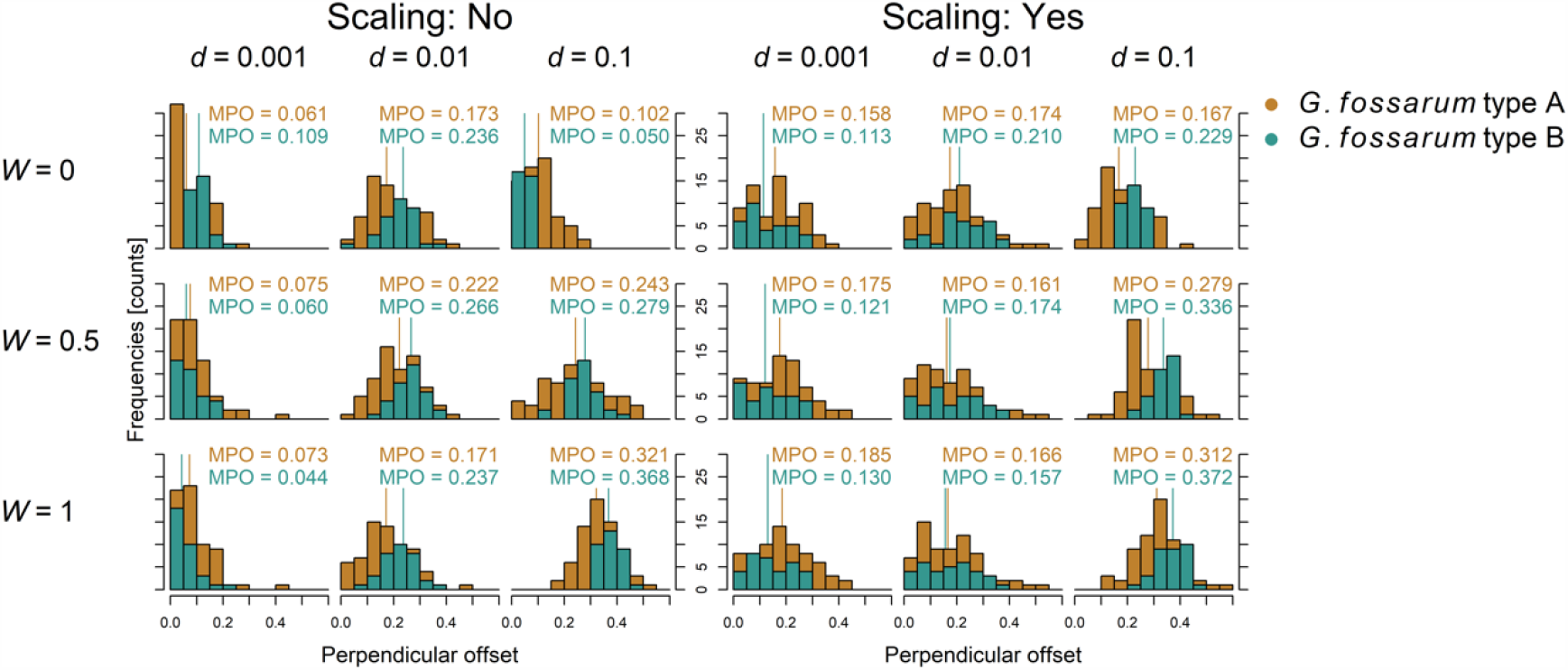
Histograms and medians of the perpendicular offsets (MPO) between all 18 stochastic simulation scenarios and the empirical mean observed heterozygosity values for both species of the *Gammarus fossarum* complex (type A: orange symbols, type B: cyan symbols). The actual perpendicular offsets of all 18 scenarios is given in Figure S7.

In addition, a qualitative, visual inspection of local polynomial regression fits of simulated and empirical data confirmed the observed large-scale pattern matches, such as higher allelic richness in downstream nodes. Generally, simulations with low dispersal rates (*d* = 0.001) fell closer to the observed range of values compared to higher dispersal rates (Figure S14 and S15). Upstream movement probability showed no consistent signal when compared to the empirical data. Unlike results from perpendicular offsets where simulations without scaled carrying capacities outperformed simulations with scaled carrying capacities, the latter successfully predicted peaks of allelic richness at the corresponding network position (e.g. *d* = 0.001, *W* = 0, scaled carrying capacity matches empirical data from *G. fossarum* type A).

## Discussion

Combining stochastic simulations and empirical data from two amphipod species within a large riverine network showed a clear signature of spatial configuration and connectivity on their genetic diversity and structure across both metapopulations. The stochastic simulations embraced the specific nature of riverine networks by using a realistic representation of the riverine network (Carraro et al., 2020) with 2,401 nodes, helping to dissect the relevant processes explaining the genetic diversity.

Past theoretical (Blanchet et al., 2020; Fronhofer & Altermatt, 2017; Morrissey & de Kerckhove, 2009; Paz-Vinas et al., 2015) and empirical (Fourtune et al., 2016; Paz-Vinas et al., 2018; Seymour et al., 2016) studies have postulated specific effects of riverine network configuration on the genetic diversity and structure of aquatic organisms. Here, we assessed empirical data across large natural metapopulations of freshwater amphipods (Figure 1), and found that measures of local genetic diversity such as allelic richness and heterozygosity were higher in more central nodes of the network (DIDG; Paz-Vinas et al., 2015). This supports theoretical expectations when the network entails some dispersal limitation. However, for the node-specific genetic measures, allelic richness was best explained by upstream distance from the outlet node and total catchment area, whereas observed heterozygosity was best explained by undirected closeness centrality and directed betweenness centrality (Figure 2). The two different measures of genetic diversity hence were best explained by different network metrics, which is surprising and implies biologically different processes or a different focus of these two measures. The upstream distance likely captures aspects of colonization legacy (either on the species itself or on some of its competitors) and may better correlates with larger biogeographic regions. This legacy seems best captured by allelic richness. Contrarily, closeness centrality better captures overall connectivity within the metapopulation, hence the more short-term effect of dispersal seems to manifest in the observed heterozygosity.

We then compared the empirical data to stochastic simulations run under different scenarios, to identify the main drivers of population genetic structuring (Figure 3). Importantly, the simulations were run on the same, and spatially realistic, graph representation of the empirical river network where the population genetics data originated from, whereas previous studies on population genetic diversity in riverine networks strongly relied on more artificial representations of riverine networks (e.g. Paz-Vinas et al., 2015). The comparison using SPO (Figure S6 and S7) and MPO (Figure 4 and 5) showed that simulations with low dispersal rate were best matching the observed patterns. This strongly suggests that the magnitude of dispersal rate has the most pronounced effect on population genetic diversity. Higher dispersal rates may homogenize populations (Bohonak, 1999). Hence, observing clear structure in the genetic diversity of a metapopulation suggest only moderate dispersal or connectivity. Our simulations required low rates of dispersal to match the empirically observed patterns (Figure S3). However, even the highest dispersal rates used in our model (*d* = 0.1) led to clear spatial structure in population genetics in combination with restricted or moderate upstream movement (Figure 3 and S2). Thus, the comparison between empirical data and simulations support the notion that riverine networks impose such strong restrictions on connectivity between nodes that spatial genetic diversity can be maintained despite high dispersal rates.

We varied upstream movement considerably in our simulations, from upstream and downstream movements being equally likely (*W* = 1) to excluding upstream movement completely (*W* = 0). This biologically meaningful change was not reflected in comparable differences in simulation matches (Figure S4). Among the best fitting simulations were simulations with an equal probability of up- and downstream movement as well as those without upstream movement. This highlights that it is the riverine network topology itself and dispersal in this very defined setting and less so the directionality of dispersal in riverine networks that is shaping population genetic structure. The riverine network imposes a low and restricted connectivity compared to a lattice-type landscape and seems to reinforce the role of dispersal, overruling the directionality. When comparing simulations with identical dispersal rates, those including restricted upstream movement (lower values of *W*) fitted slightly better than simulations with no movement directionality, indicating some influence of asymmetric gene flow in generating the observed patterns (Fraser et al., 2004). With low dispersal rates, the upstream movement probability did not play a major role in improving the model fit. However, completely asymmetric gene flow seems unrealistic, given the subtle differences within simulations using the same dispersal rate (Morrissey & de Kerckhove, 2009). One major unknown, requiring further empirical studies, is how mobile the studied species actually are; often, they are considered comparably poor dispersers (Elliott, 2003; Weiss & Leese, 2016) or mostly transported passively by drift, while some studies suggest them being rather mobile (Meijering, 1972; Žganec et al., 2013). Our results do not allow us to draw a conclusion regarding the process that drives upstream movement, and how much of the dispersal is active versus passive (e.g., downstream transport or dislocation by vectors). Possibly, our view of the riverine network being downstream oriented is not what the studied amphipods experience, and their benthic life-style may mitigate downstream flow considerably (Statzner & Holm, 1989).

Surprisingly, scaling the carrying capacity of nodes with the square-root of the total catchment area (Ozerov et al., 2012), and thus making the habitat capacity more realistic, lowered simulation fits to empirical data when compared to the unscaled counterparts in two thirds of the simulation cases, irrespective of the species (Figure S5). The scaling generally increased the range of the response values, making SPO and MPO increase. In the empirical data, however, we did not find comparably high levels of allelic richness and observed heterozygosity. We cannot exclude that the chosen scaling function does not correspond to the realized distribution of carrying capacities nor that the carrying capacity does not scale at all. Long-term data from the lower part of the Rhine in Switzerland, that is, in a very large stream, indicate very high densities of *G. fossarum* species compared to commonly observed densities in upstream reaches (Mürle et al., 2008), contradicting the non-scaling of carrying capacity. In addition, *G. fossarum* type B is a slightly more tolerant species and can also be found in anthropogenically more affected streams, while *G. fossarum* type A is the typical amphipod species of near-natural headwater streams (Eisenring et al., 2016). Hence, we expected *G. fossarum* type B to be more common in larger streams (i.e., larger total catchment area) and therefore showing less variance in occupied carrying capacities. However, a simple two-sample Kolmogorov-Smirnov test indicated that the samples we had at hand did not differ significantly in their total catchment area and consequently their stream width (data not shown). This could explain that the model fits were worse in both *G. fossarum* type A and type B when scaling the carrying capacity.

The comparisons of local polynomial regression fits between empirical and simulated data showed some remarkable correspondence (Fig. S14 and S15). Observed peaks of allelic richness were also present at the corresponding network positions in some of the stochastic simulations whereas they were completely absent in other simulations. Some parameter combinations showed opposite patterns to the empirically observed data. This highlights that even simulations with only few assumptions and without any environmental influence or biotic interactions successfully predict large-scale population genetic patterns in a riverine network. Again, connectivity of nodes and their carrying capacities coupled with organismal dispersal can drive empirically observed patterns of the genetic diversity of purely aquatic organisms.

In conclusion, our study showed a pronounced effect of dispersal rate and the riverine network on the genetic diversity and structure of two amphipod species across a large spatial extent. In comparison to dispersal rate, the directionality of dispersal was of minor importance, at least for the species studied. Furthermore, assumed increasing local population sizes with increasing downstream distance were not important in shaping population genetic diversity. This indicates that for understanding and protecting genetic diversity of riverine organisms, the network connectivity is the single most important aspect to be considered.

## Acknowledgements

We thank Pravin Ganesanandamoorthy, Elvira Mächler, Katri Seppälä, and Anja Marie Westram for help during the laboratory work. Jukka Jokela, Anja Marie Westram and Luca Carraro provided helpful feedback on the manuscript and Felix Moerman helped with running the simulations. Funding is from the Swiss National Science Foundation Grant No PP00P3_150698 (to F.A.). This is publication ISEM-2021-YYY of the Institut des Sciences de l’Evolution – Montpellier.

## Data Accessibility

The data are deposited on GitHub under the following digital object identifier 10.5281/zenodo.4321240: Microsatellite data (Gammarus_all_microsat_data_Rhein_2018.txt); Graph object (Gfos_Network.RData); Simulation code (pop_gen_gammaridae_v0.cpp); Analysis script for R (PopGenNet20201210.R).

## Author contributions

FA and EAF designed the research. FA, EAF, RA performed coding and planning of molecular work. EAF and RA analysed data. RA and FA wrote the first draft of the paper. All authors commented on the final draft.

