## Supplementary material for "Dispersal rate and riverine network connectivity shape the genetic diversity of freshwater amphipod metapopulations": S1

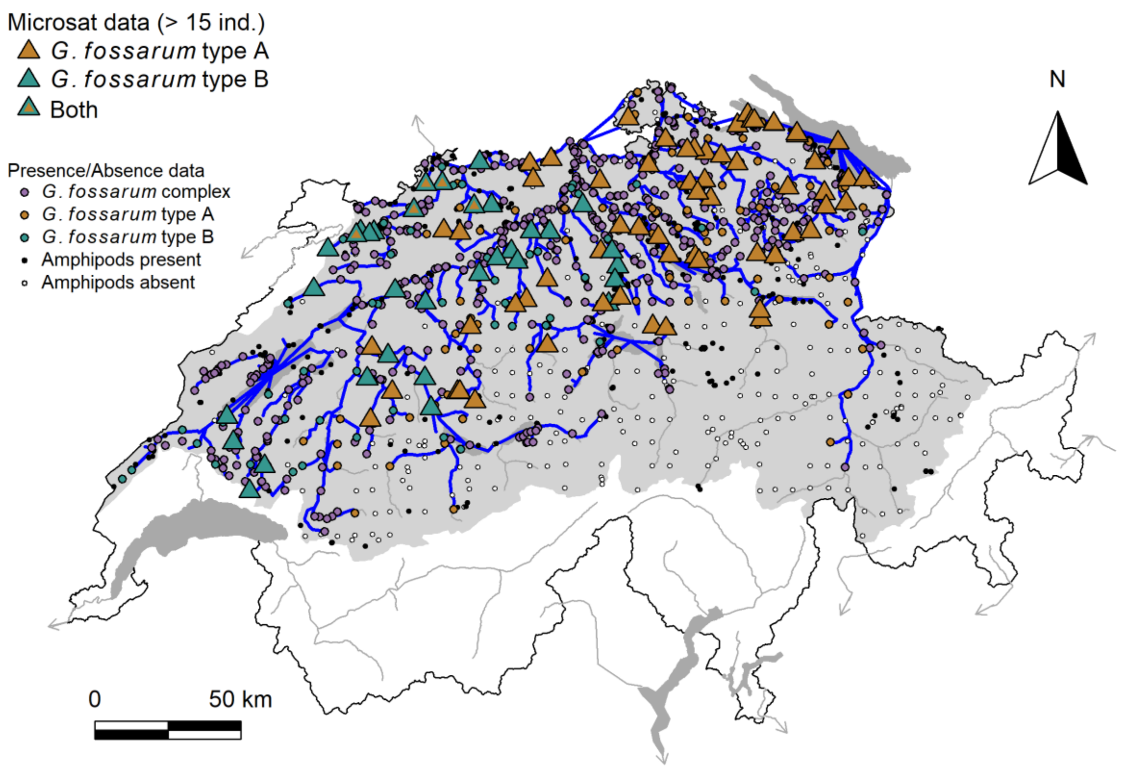


**Figure S1** Graph representation of the riverine network (blue lines) that is accessible to *Gammarus fossarum* type A and type B within the total Rhine catchment (shading). It is based on a 2 $km^2$ subcatchment representation of streams and rivers of Switzerland. Geodata source: Federal Office of Topography & Federal Office for the Environment
