## Supplementary material for "Dispersal rate and riverine network connectivity shape the genetic diversity of freshwater amphipod metapopulations": S2

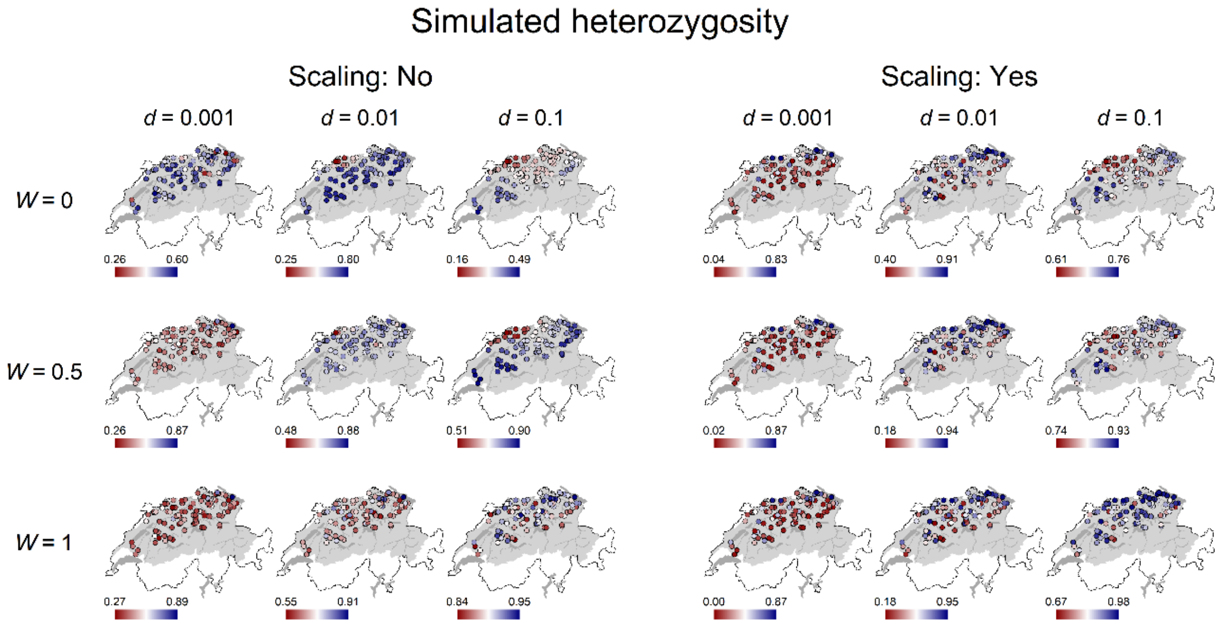


**Figure S2** Maps depicting the predicted heterozygosity for all 18 stochastic simulation scenarios show different spatial structuring along the Rhine riverine network of Switzerland. The gradient legends show observed heterozygosity. Their ranges are adjusted for each map for the best visual representation of spatial structuring. Geodata source: Federal Office of Topography & Federal Office for the Environment
