## Supplementary material for "Dispersal rate and riverine network connectivity shape the genetic diversity of freshwater amphipod metapopulations": S3

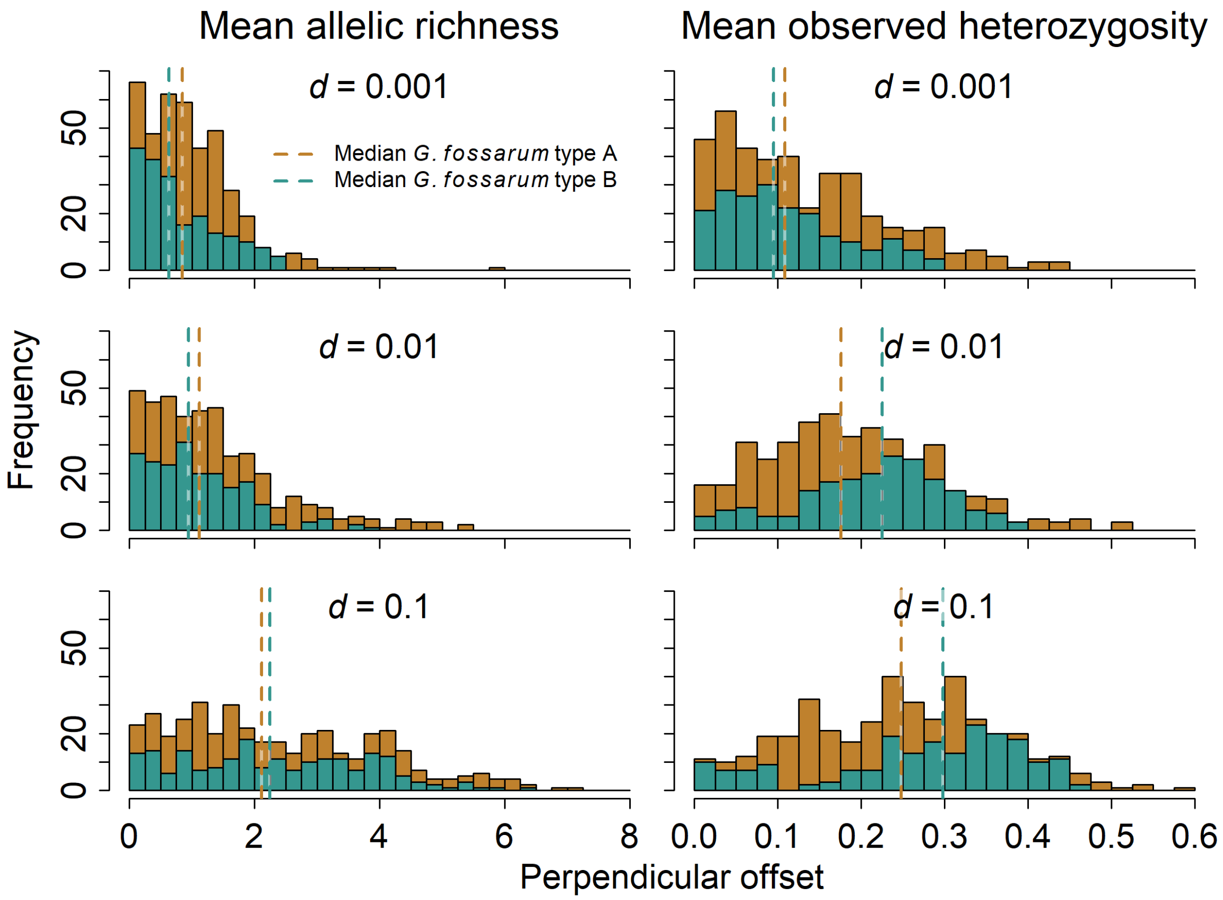


**Figure S3** Different dispersal rates in the simulations resulted in strong variability on the simulation fit, reflected in perpendicular offset differences. Simulations with low dispersal rates ($d = 0.001$) were superior to simulations with higher dispersal rates. High dispersal rates generally resulted in the worst model fits.
