## Supplementary material for "Dispersal rate and riverine network connectivity shape the genetic diversity of freshwater amphipod metapopulations": S4

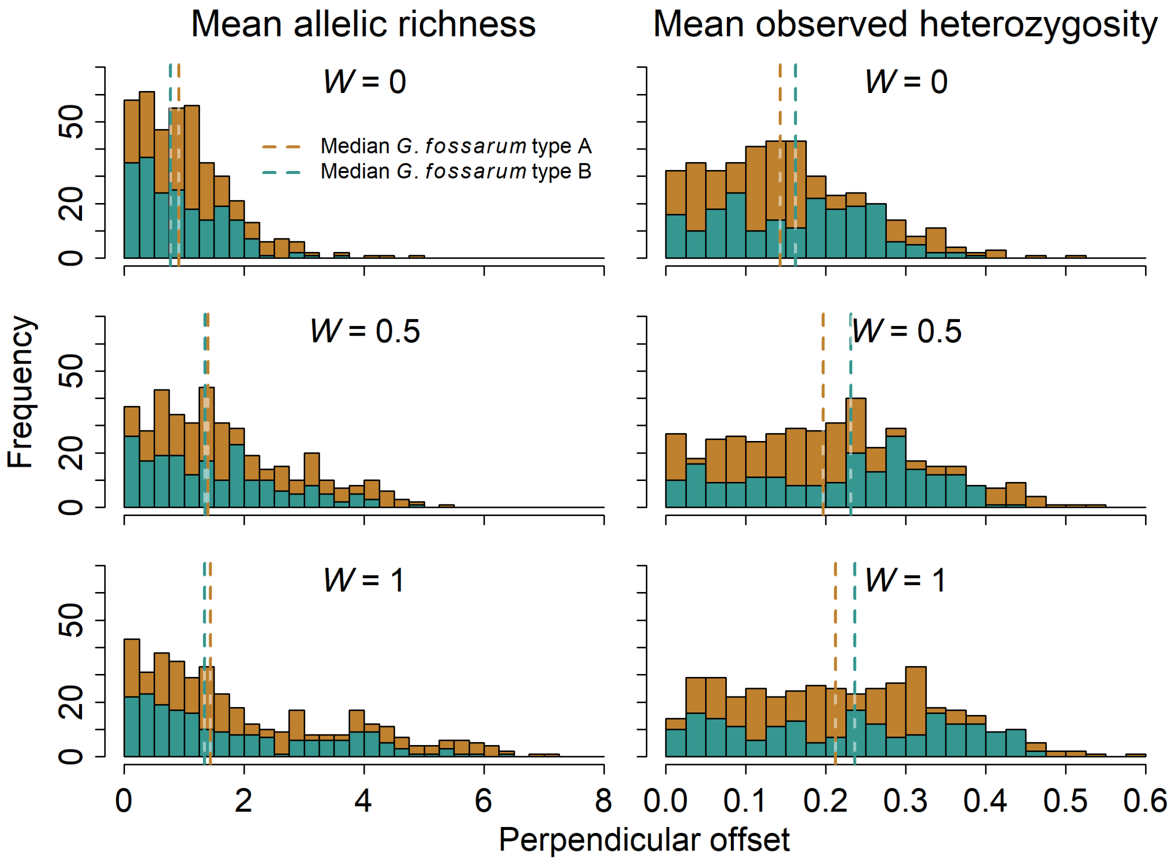


**Figure S4** Changing upstream movement probabilities in the simulations resulted in slight shifts of model fits to empirical data in comparison to dispersal rate (see Fig. S3). The clearest signal results when allowing for upstream dispersal (*W* = 0.5 and *W* = 1), weakening model fits as reflected in perpendicular offset differences. Simulations with no upstream dispersal (*W* = 0) were superior to those simulations.
