## Supplementary material for "Dispersal rate and riverine network connectivity shape the genetic diversity of freshwater amphipod metapopulations": S5

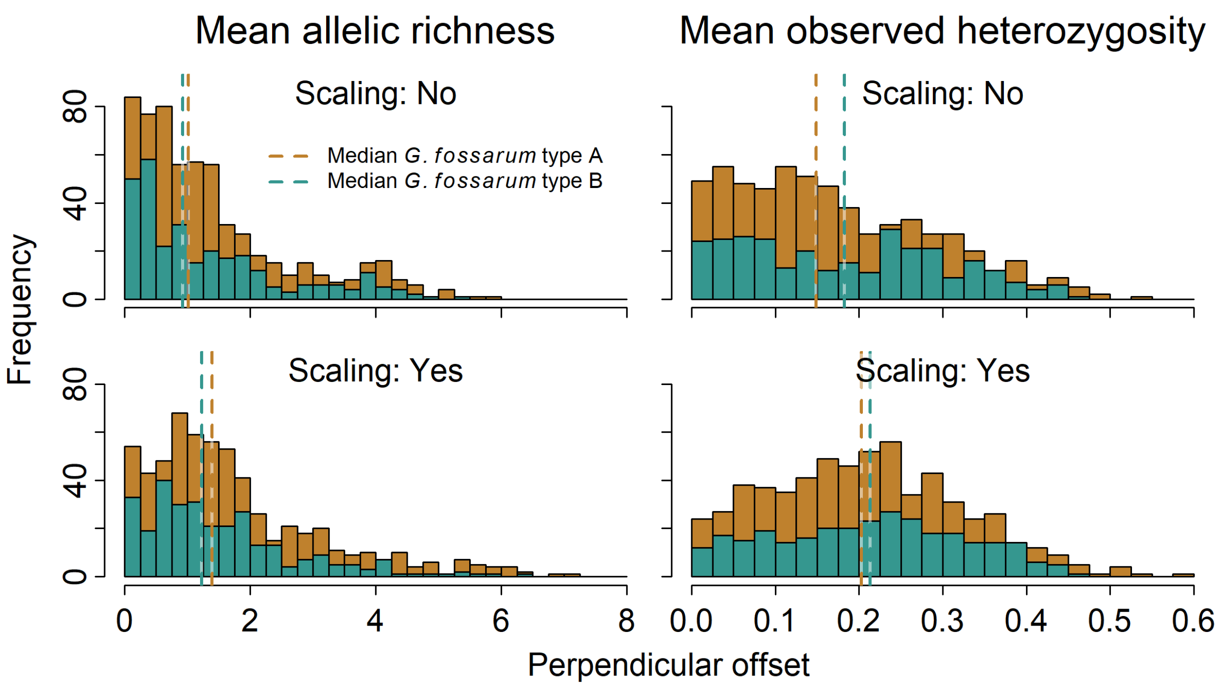


**Figure S5** Scaling the habitat carrying capacity showed the lowest impact on the variability of the response variable. Generally, simulations without scaling of the carrying capacity ($K = 0$) outperformed the ones where carrying capacity scaled with the square-root of the total catchment area.
