## Supplementary material for "Dispersal rate and riverine network connectivity shape the genetic diversity of freshwater amphipod metapopulations": S7

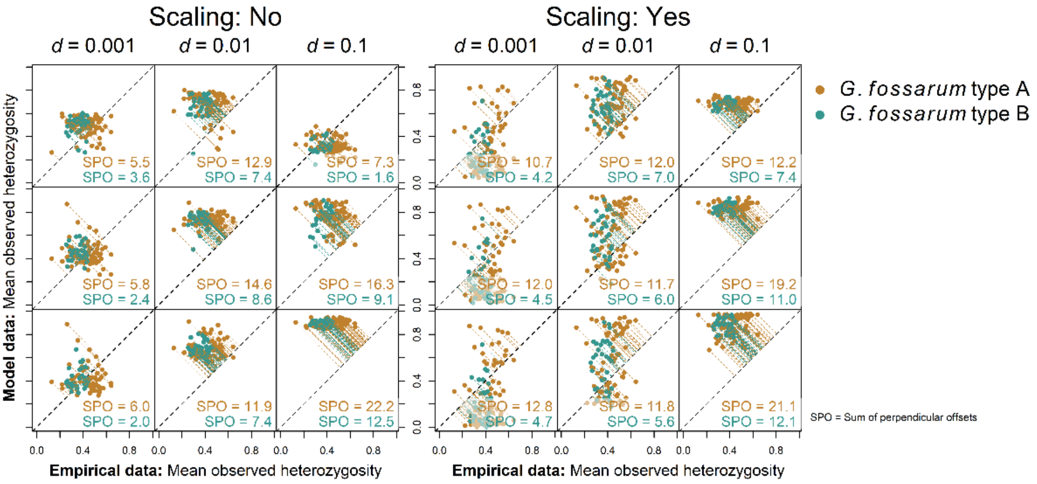


**Figure S7** Perpendicular offsets between observed heterozygosity value pairs of all 18 stochastic simulation scenarios and the empirical data. The sum of the perpendicular offsets (SPO) served as a goodness-of-fit measure. SPO takes into account the overall spread of simulated values from their empirical counterpart, where larger SPO indicates a poorer fit.
