## Supplementary material for "Dispersal rate and riverine network connectivity shape the genetic diversity of freshwater amphipod metapopulations": S11

**Table S11** Ranking of the 18 stochastic simulations for their fit to the empirically assessed mean observed heterozygosity values according to their median of perpendicular offsets (MPO). Listed are the value for MPO and the corresponding varying parameter values (dispersal rate *d*, upstream movement probability *W*, and scaling of habitat carrying capacity *K*).

| Rank | Mean observed heterozygosity | | | | | | | |
|---|---|---|---|---|---|---|---|---|
| | *G. fossarum* type A | | | | *G. fossarum* type B | | | |
| | MPO | *d* | *W* | *K* | MPO | *d* | *W* | *K* |
| 1 | 0.061 | 0.001 | 0 | 0 | 0.044 | 0.001 | 1 | 0 |
| 2 | 0.073 | 0.001 | 1 | 0 | 0.050 | 0.1 | 0 | 0 |
| 3 | 0.075 | 0.001 | 0.5 | 0 | 0.060 | 0.001 | 0.5 | 0 |
| 4 | 0.102 | 0.1 | 0 | 0 | 0.109 | 0.001 | 0 | 0 |
| 5 | 0.158 | 0.001 | 0 | 1 | 0.113 | 0.001 | 0 | 1 |
| 6 | 0.161 | 0.01 | 0.5 | 1 | 0.121 | 0.001 | 0.5 | 1 |
| 7 | 0.166 | 0.01 | 1 | 1 | 0.130 | 0.001 | 1 | 1 |
| 8 | 0.167 | 0.1 | 0 | 1 | 0.157 | 0.01 | 1 | 1 |
| 9 | 0.171 | 0.01 | 1 | 0 | 0.174 | 0.01 | 0.5 | 1 |
| 10 | 0.173 | 0.01 | 0 | 0 | 0.210 | 0.01 | 0 | 1 |
| 11 | 0.174 | 0.01 | 0 | 1 | 0.229 | 0.1 | 0 | 1 |
| 12 | 0.175 | 0.001 | 0.5 | 1 | 0.236 | 0.01 | 0 | 0 |
| 13 | 0.185 | 0.001 | 1 | 1 | 0.237 | 0.01 | 1 | 0 |
| 14 | 0.222 | 0.01 | 0.5 | 0 | 0.266 | 0.01 | 0.5 | 0 |
| 15 | 0.243 | 0.1 | 0.5 | 0 | 0.279 | 0.1 | 0.5 | 0 |
| 16 | 0.279 | 0.1 | 0.5 | 1 | 0.336 | 0.1 | 0.5 | 1 |
| 17 | 0.312 | 0.1 | 1 | 1 | 0.368 | 0.1 | 1 | 0 |
| 18 | 0.321 | 0.1 | 1 | 0 | 0.372 | 0.1 | 1 | 1 |
