## Supplementary material for "Dispersal rate and riverine network connectivity shape the genetic diversity of freshwater amphipod metapopulations": S13

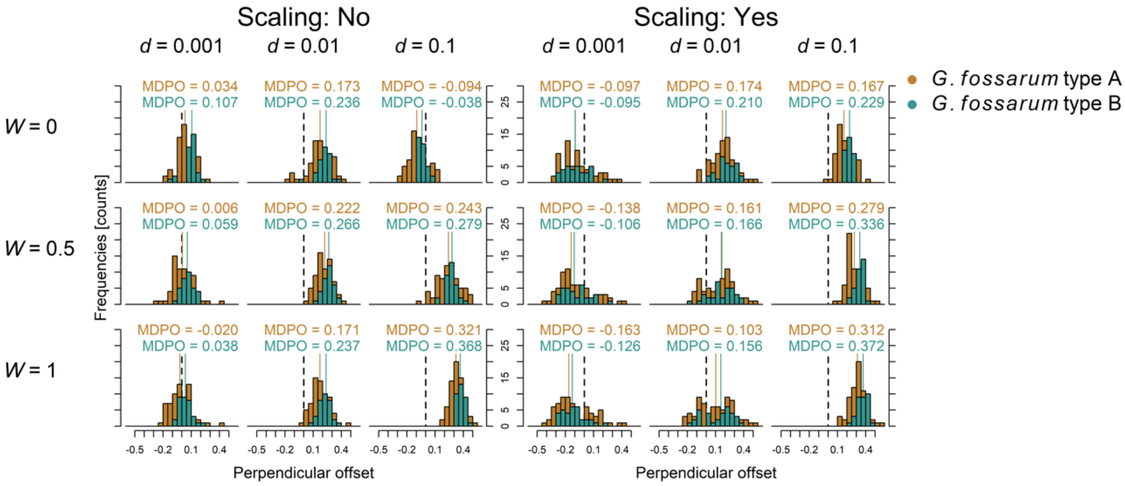


**Figure S13** Histograms and medians of the directed perpendicular offsets (MDPO) between all 18 stochastic simulation scenarios and the empirically assessed mean observed heterozygosity values. The directed perpendicular offset does take into account if points are above or below the vertical (1:1) line. The dotted vertical line shows zero offset, corresponding to a perfect fit between simulation and empirical data.
