## Supplementary material for "Dispersal rate and riverine network connectivity shape the genetic diversity of freshwater amphipod metapopulations": S14

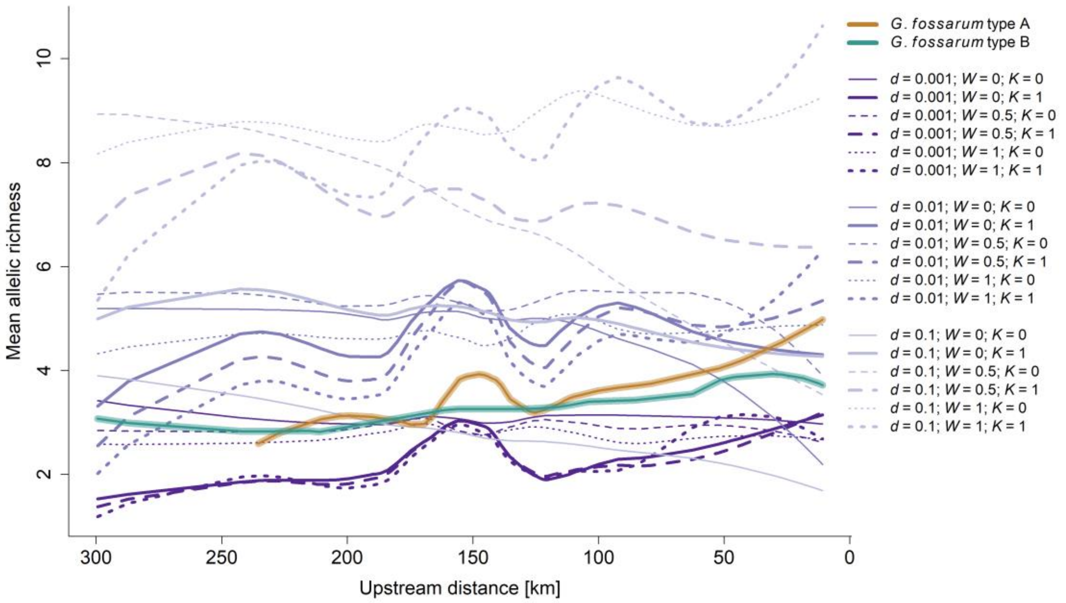


**Figure S14** Empirically assessed mean allelic richness of *G. fossarum* type A and type B in comparison to all the 18 simulated scenarios. The data are shown in relation to upstream distance from the outlet node within the riverine network. The lines correspond to the local polynomial regression fit.
